# Primed bovine embryonic stem cell lines can be derived at diverse stages of blastocyst development with similar efficiency and molecular characteristics

**DOI:** 10.1101/2024.11.15.623897

**Authors:** Carly Guiltinan, Ramon C. Botigelli, Juliana I. Candelaria, Justin M. Smith, Rachel B. Arcanjo, Anna C. Denicol

**Author notes:** Current affiliation: Department of Biomolecular Engineering, University of California, Santa Cruz, Santa Cruz, CA, USA. Current affiliation: Department of Biochemistry and Molecular Biology, Uniformed Services University of the Health Sciences, Bethesda, MD, USA. Current affiliation: Departamento de Genética e Evolução, Universidade Federal de São Carlos, São Carlos, SP, Brazil. Corresponding author: Anna C. Denicol, Associate Professor – Department of Animal Science University of California, Davis 450 Bioletti Way, Davis, CA 95616, USA.

## Abstract

In this study, we established bovine embryonic stem cell (bESC) lines from early (eBL) and full (BL) blastocysts to determine the efficiency of bESC derivation from an earlier embryonic stage and compare the characteristics of the resulting lines. Using established medium and protocols for derivation of primed bESCs from expanded blastocysts, we derived bESC lines from eBLs and BLs with the same efficiency (4/12 each, 33%). Regardless of original blastocyst stage, bESC lines had a similar phenotype, including differentiation capacity, stable karyotype, and pluripotency marker expression over feeder-free transition and long-term culture. Transcriptome and functional analyses indicated that eBL– and BL-derived lines were in primed pluripotency. We additionally compared RNA-sequencing data from our lines to bovine embryos and stem cells from other recent reports, finding that base medium was the predominant source of variation among cell lines. In conclusion, our results show that indistinguishable bESC lines can be readily derived from eBL and BL, widening the pool of embryos available for bESC establishment. Finally, our investigation points to sources of variation in cell phenotype among recently reported bESC conditions, opening the door to future studies investigating the impact of factors aside from signaling molecules on ESC derivation, maintenance, and performance.

**SUMMARY STATEMENT:** Primed bovine embryonic stem cell lines can be readily established from early blastocysts and are indistinguishable from full blastocyst-derived lines, widening the pool of embryos available for stem cell derivation.

## INTRODUCTION

Embryonic stem cells (ESCs) are pluripotent stem cells (PSCs) that are stabilized in vitro from the epiblast of the inner cell mass (ICM) of a blastocyst. In vivo, the epiblast is only pluripotent transiently, first becoming fate-restricted and then gastrulating to form the three embryonic germ layers – which eventually differentiate into all somatic cell types of the body – and the germline. Modulation of key signaling pathways has enabled long-term culture of ESC lines in an undifferentiated, self-renewing state in several mammalian species (Alberio et al., 2010; Hildebrandt et al., 2018; Kaufman and Evans, 1981; Martin, 1981; Thomson, 1998; Thomson et al., 1995). After many years of collective effort, conditions that support the establishment of stable bovine ESC (bESC) lines were described in 2018 (Bogliotti et al., 2018).

Previous work has shown that distinct states of pluripotency – which resemble the developmental states of the epiblast from blastocyst to gastrula – can be captured in vitro by modulation of signaling factors in the medium. The naïve and primed states flank the pluripotent continuum, while intermediate states including formative pluripotency have recently been reported (Kinoshita et al., 2021a; Nichols and Smith, 2009; Smith, 2017; Yu et al., 2021). Formative PSCs resemble the peri-gastrulation epiblast, at which time the cells of the epiblast are preparing for multi-lineage competence but not yet biased toward a specific cell lineage (Smith, 2017). As a result, these cells are ideal for directed in vitro differentiation toward derivatives of the three germ layers. Notably, the formative interval of development is also the time at which primordial germ cells (PGCs) are specified in the epiblast, and therefore formative PSCs are permissive to efficient direct induction of PGC-like cells, unlike naïve or primed PSCs which must go through intermediate pre-differentiation steps (Hayashi et al., 2011; Sasaki et al., 2015). Expanded potential stem cells (EPSCs) have also been recently described, which have the capacity to produce derivatives of both embryonic and extraembryonic lineages when integrated into chimeras (Gao et al., 2019; Yang et al., 2017; Zhao et al., 2021). In addition to distinctions in the culture medium, derivation of mouse EPSC lines was performed by isolating individual blastomeres of 8-cell embryos, unlike ESCs which are established from blastocysts, indicating the potential of harboring less advanced states of pluripotency by targeting earlier embryonic stages.

Following the initial report, alternative approaches to stabilize PSCs from cattle have recently been described. For instance, modification of the base medium to a commercially available product, use of fully defined media components, and adaptation to feeder-free matrices has made bESC culture more scalable (Kinoshita et al., 2021b; Soto et al., 2021). Bovine EPSCs have also been reported, which have distinct transcriptomic and epigenetic features from primed bESCs (Zhao et al., 2021). A recent report targeted the epiblast of bilaminar disc-staged day-11 porcine embryos for establishment of primed embryonic disc stem cells (EDSCs) in fully defined conditions, indicating that a wide range of epiblast stages could be used for the establishment of PSC lines in livestock (Kinoshita et al., 2021b). This report utilized the same culture condition to produce bovine PSCs from day 7-9 blastocysts, but did not compare the resulting cell lines based on the embryo stage from which they were derived. Characterization of the first reported bovine ESC and EDSC lines has indicated that these cells are in primed pluripotency, which limits their applications due to fate-biasedness compared to cells in a naïve or formative pluripotent state. On the other hand, some groups have demonstrated the capacity of bovine induced PSC (iPSC) lines to be stabilized in a naïve-like state (Pillai et al., 2021; Su et al., 2021), although this has not yet been achieved for bESCs.

The objective of this study was to assess the effect of the embryonic stage from which bESC lines were derived when cells were otherwise established using the same methods and maintained in the same conditions. Previously, bESC lines were derived from full, expanded, or hatched blastocysts, but the embryo stage was not kept track of when assessing cell lines (Bogliotti et al., 2018; Kinoshita et al., 2021b; Shirasawa et al., 2024; Soto et al., 2021). We first set to determine if bESC lines could be established from early blastocysts (eBL), and then to assess how the resulting cell lines compare to those established from full, non-expanded blastocysts (BL). Such knowledge would indicate whether a wider range of bovine embryos could be used for bESC derivation and if certain embryonic stages could be targeted to give specific features to bESC lines. Additionally, we sought to provide a broad transcriptomic comparison of bovine ESC, EDSC, and EPSC lines from recent reports to provide context of the distinct features of these cells based on the alternative strategies to establish lines.

## RESULTS

### Stable, pluripotent bESC lines can be established from early blastocysts with similar efficiency to full blastocysts

Bovine ESC lines could be established from eBL with the same efficiency as BL (4/12 embryos, 33.0%) using described derivation methods in NBFR medium(Soto et al., 2021;Fig. 1A-B). The resulting lines homogenously expressed the core pluripotency proteins, OCT4, SOX2 (Fig. 1C), and NANOG (Fig. S1A), which was maintained upon transition to feeder-free culture on VTN-N despite changes in cell morphology from colonies to monolayer-like growth (Fig. 1C), as has been described for this condition (Soto et al., 2021). Of the seven lines that were tested, six were determined to be female (85%; Fig. S1B). Two female cell lines derived from each blastocyst stage were maintained on VTN-N for more than 30 passages, while retaining stable karyotype (2n = 60; Fig. 1D). By the time of writing this manuscript, we have extensively cultured the resulting cell lines from this experiment for over 3 years with stable phenotype. We also assessed in vivo differentiation capacity by teratoma formation (Hentze et al., 2009), and found that bESC lines derived from both blastocyst stages were capable of giving rise to all three embryonic germ layers (Fig. 1E).

**Figure 1.**
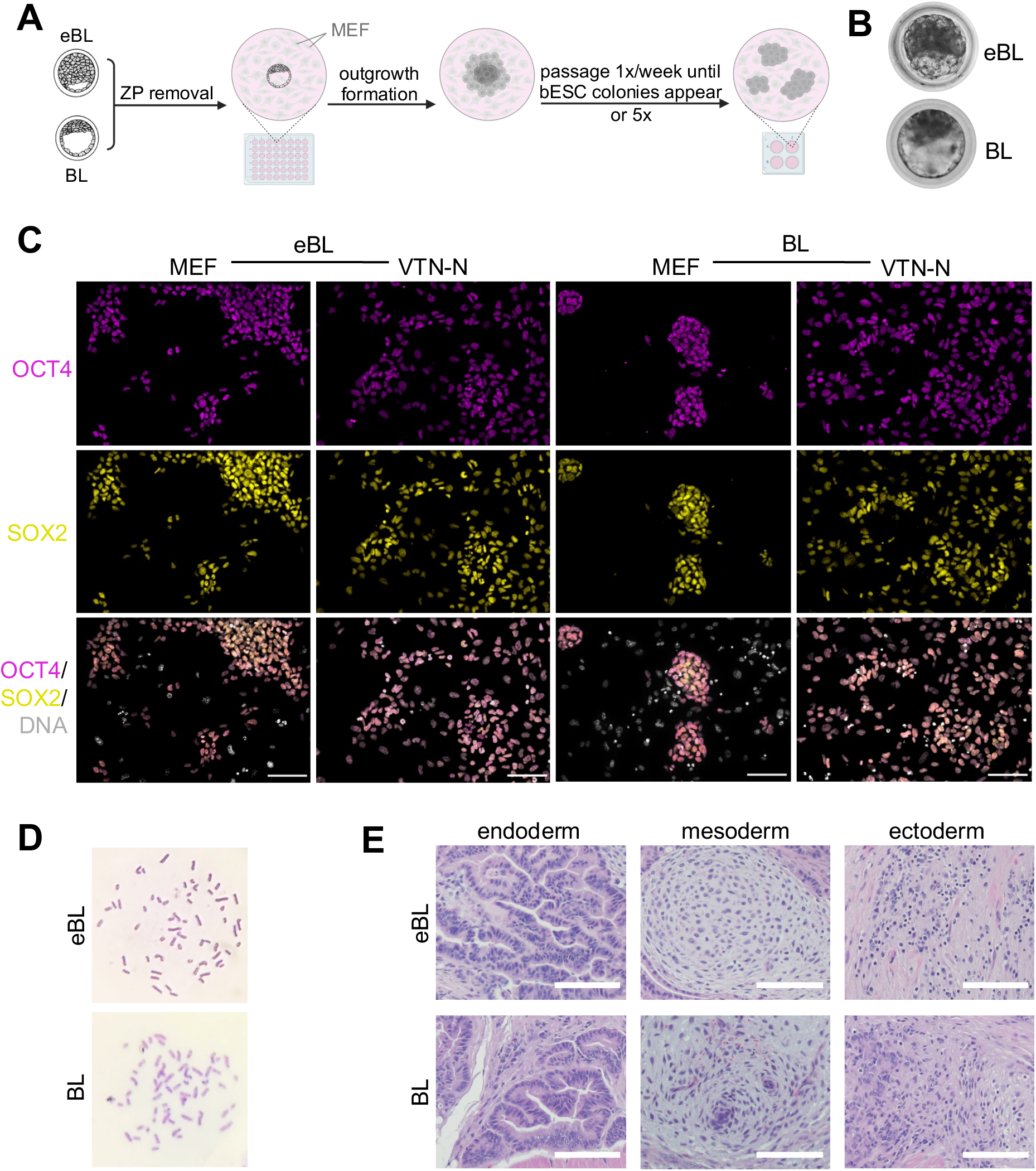
Stable, pluripotent bovine embryonic stem cells can be established from early and full blastocyst-stage embryos. **(A)** Schematic showing bovine ESC derivation methods. ZP = zona pellucida; MEF = mouse embryonic fibroblasts. **(B)** Representative images of early (eBL) and full (BL) blastocysts used for bESC derivation. **(C)** Representative immunofluorescence images of bESCs derived from eBL and BL on MEF feeders and feeder-free on vitronectin (VTN-N) at passage 14. Magenta indicates OCT4; yellow indicates SOX2; gray indicates DNA. Scale bar, 100 μm. **(D)** Karyotype analysis of bESC lines from eBL and BL at passage 30 by Giemsa chromosome staining. **(E)** Teratoma formation from eBL and BL-derived lines with differentiation into endoderm-derived tissue (gut epithelium), mesoderm-derived tissue (cartilage, mucinous mass), and ectoderm derived tissue (neuronal epithelium). Scale bar, 100 μm.

### Bovine ESC lines are similar and exhibit features of primed pluripotency when cultured in the same conditions, regardless of initial blastocyst stage

Upon seeing little difference in bESC lines through the initial assessments, we performed global RNA sequencing of two female bESC lines from each embryonic stage at passage 9 to investigate potential differences in global transcriptome. We found that eBL– and BL-derived bESC lines had subtle differential gene expression, including 5 upregulated and 22 downregulated genes (Fig. 2A), which likely can be attributed to line-to-line heterogeneity that is universally common for ESCs. Moreover, markers associated with pluripotency and primed progression were not differentially expressed among the lines based on embryo stage (Fig. 2B). All bESC lines had lower relative expression of naïve pluripotency markers (e.g., *KLF4*, *ESRRB*, *TBX3*, *DPPA3, MYC*) and elevated markers of primed progression (e.g., *OTX2*, *ZIC2*, *XIST*, *HDAC2*, *DNMT3B*) but not early lineage commitment (e.g., *EOMES*, *MIXL1*). These results were consistent with prior findings that bESCs cultured with the same cytokines were in a primed state of pluripotency (Bogliotti et al., 2018). We additionally integrated these samples with RNA-sequencing data of the four lines after feeder-free transition and extended culture (passage 23-32), as well as epiblast samples from bovine embryos at day 7, 14, and 17 of development (Bernardo et al., 2018; Fig. 2C). This showed that the pattern of differential gene expression among the cell lines based on embryo stage of origin was not maintained long-term, and suggest that extended culture and matrix affect bESC transcriptome more than the initial embryo stage does. In addition, this analysis showed that bESCs in NBFR medium resembled the epiblast at a stage intermediate to day 7 (blastocyst) and 17 (peri-gastrula), with more similarity to day 14-17 vs. day 7 epiblasts, which is consistent with a primed pluripotent state (Nichols and Smith, 2009). Collectively, these findings show that the blastocyst stage from which ESCs are derived does not significantly alter the transcriptome of the resulting cell lines by 9 passages in the same condition.

**Figure 2.**
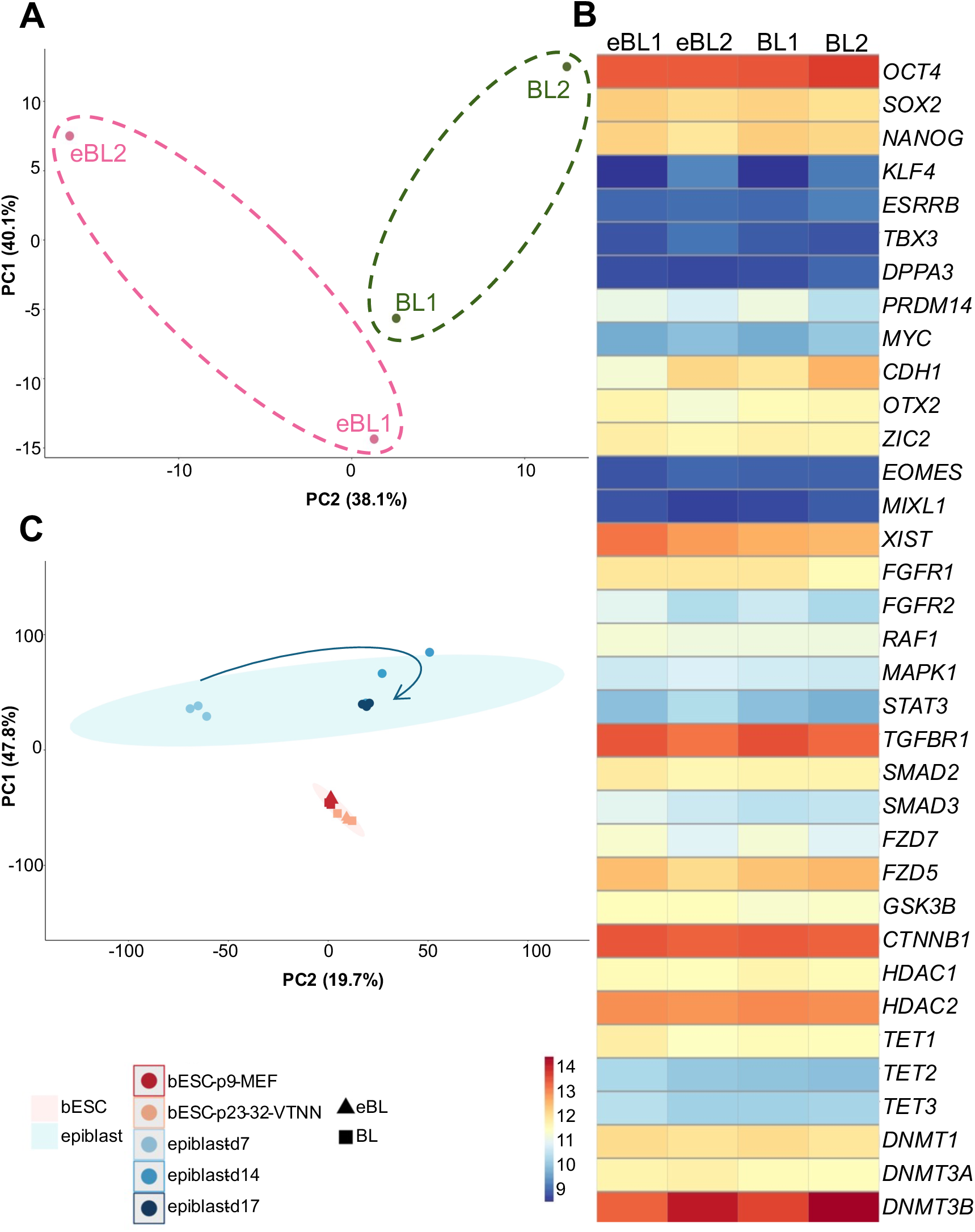
Bovine ESC lines are transcriptomically similar regardless of blastocyst stage from which they were derived. **(A)** Principal component analysis from RNA-sequencing of bovine ESC lines derived from early (eBL) and full (BL) blastocysts at passage 9 on mouse embryonic fibroblast (MEF) feeders. **(B)** Heatmap of genes associated with pluripotency, primed progression, and epigenetic status for eBL– and BL-lines at passage 9 on MEFs. **(C)** Principal component analysis from integrated analysis of RNA-sequencing of bESC lines from eBL and BL on MEFs at passage 9, on vitronectin (VTN-N) at passage 23-32, and bovine epiblast at day 7, 14, and 17 of embryonic development. Arrow indicates developmental progression of the epiblast.

We also sought to functionally assess the pluripotent state of cell lines. We first tested the dependency of our cell lines on exogenous FGF2. All reports of primed bESCs rely upon inclusion of FGF2 in the medium (Bogliotti et al., 2018; Kinoshita et al., 2021b; Li et al., 2023; Soto et al., 2021; Xiao et al., 2021), while expanded potential cells could be maintained without this factor (Zhao et al., 2021). Moreover, human formative PSCs are maintained in the absence of FGF, and primed human ESC lines show the plasticity to be transitioned to this condition (Kinoshita et al., 2021a). We found that bESCs from both blastocyst stages died within four days of FGF2 removal from the medium (Fig. 3A). Interestingly, bESCs could be stably maintained in lower FGF2 concentrations than the initial NBFR report, including as low as 5 ng/mL, without marked differences in proliferation rate over 4 days (Fig. 3A). Furthermore, our findings suggest that culture in lower exogenous FGF concentrations does not decrease pluripotency (*OCT4*, *SOX2*, *NANOG*) or drive primed progression or differentiation (*XIST*, *PAX6*; Fig. 3B). Prior work has demonstrated that the naïve to primed transition in female epiblast/PSCs is accompanied by inactivation of one X chromosome for dosage compensation during development (Heard, 2004). Since all cell lines analyzed in this experiment were female, we sought to determine the X-chromosome status by immunolocalization of H3K27me3, which labels Barr bodies (i.e., inactivated X chromosomes), and H3K9me2, which is enriched in primed ESCs. We found that regardless of the initial blastocyst stage, cell lines homogeneously had an inactivated X chromosome, consistent with a primed state (Nichols and Smith, 2009; Fig. 3C). This is in contrast to naïve cells, in which both X chromosomes are active (Nichols and Smith, 2009), and formative PSCs, which have both/heterogeneous X chromosome activity (Kinoshita et al., 2021a; Yu et al., 2021). Taken together, these analyses corroborate that bESCs derived from early and full blastocysts are in a primed state when maintained in NBFR medium.

**Figure 3.**
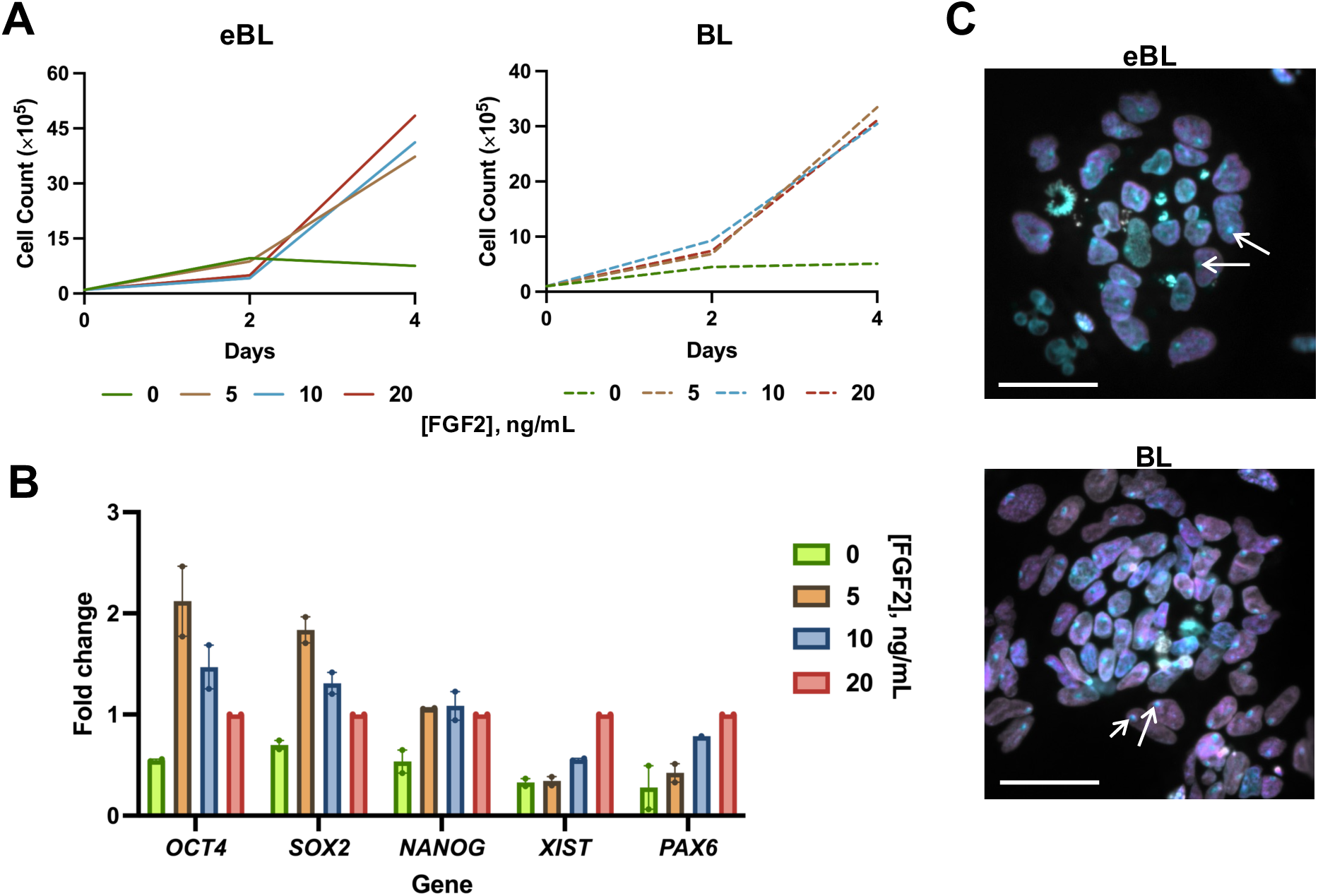
Bovine ESCs derived from early and full blastocysts show functional features of primed pluripotency. **(A)** Cell counts from NBFR-bESCs grown in typical concentration of FGF2 (20 ng/mL) or reduced concentrations of 10, 5, and 0 ng/mL FGF2. Cell counts were taken on days 2 and 4, after seeding 100,000 cells/well in respective medium on day 0. Left shows values from an early blastocyst (eBL)-derived line, right shows values from a full blastocyst (BL)-derived line. **(B)** Quantitative reverse transcriptase-PCR analysis of NBFR-bESCs after four days of culture in 0, 5, 10, or 20 ng/mL FGF2 for markers of pluripotency (*OCT4*, *SOX2*, *NANOG*), X-chromosome inactivation (*XIST*), and ectoderm lineage (*PAX6*). Analysis includes one eBL– and one BL-derived cell line. Error bars show mean with SEM, with each circle representing an experimental replicate. **(C)** Respective overlay image from immunofluorescent staining of NBFR-bESCs derived from eBL (top) and BL (bottom) at passage 15-16 for H3K27me3 (cyan), H3K9me2 (magenta), and DAPI (gray). White arrows show examples of Barr bodies, indicating an inactive X chromosome. Scale bar, 50 μm.

### Comparative RNA-sequencing analysis of bovine embryonic stem cell lines reveals potential sources of variation in cell line characteristics

Since the initial establishment of long-term stable bESC lines in 2018, rapid progress has been made by groups across the globe toward establishing alternative conditions (Bogliotti et al., 2018; Kinoshita et al., 2021b; Shirasawa et al., 2024; Soto et al., 2021; Zhao et al., 2021). We performed a comprehensive analysis of our cell lines alongside publicly-available bulk RNA-seq datasets from recently reported bovine ESC, EPSC, and EDSC lines (Bogliotti et al., 2018; Kinoshita et al., 2021b; Li et al., 2023; Zhao et al., 2021; Table 1). Interestingly, we found that the base medium was the leading source of variation among cell lines (35%; Fig. 4A). However, this variation was not highly attributable to different levels of expression of pluripotency genes (Fig. 4B). We next examined differential gene expression based on base medium by comparing similar conditions with only a different base, including mTeSR (Li et al., 2023) and our cells in N2B27 with the same factors supplemented (Fig. 4C). We identified a total of 400 upregulated and 812 downregulated genes for bESCs in N2B27 compared to mTeSR. Gene ontology (GO) analysis showed that the top upregulated biological processes of bESCs in N2B27 base medium related to metabolism (e.g., carbohydrate metabolism and biosynthesis, ribonucleotide metabolism), while those in mTeSR had positive regulation of cell communication and signal transduction compared to N2B27-cultured cells (Fig. 4C). Full GO results for enriched biological processes are shown in Table S3.

**Figure 4.**
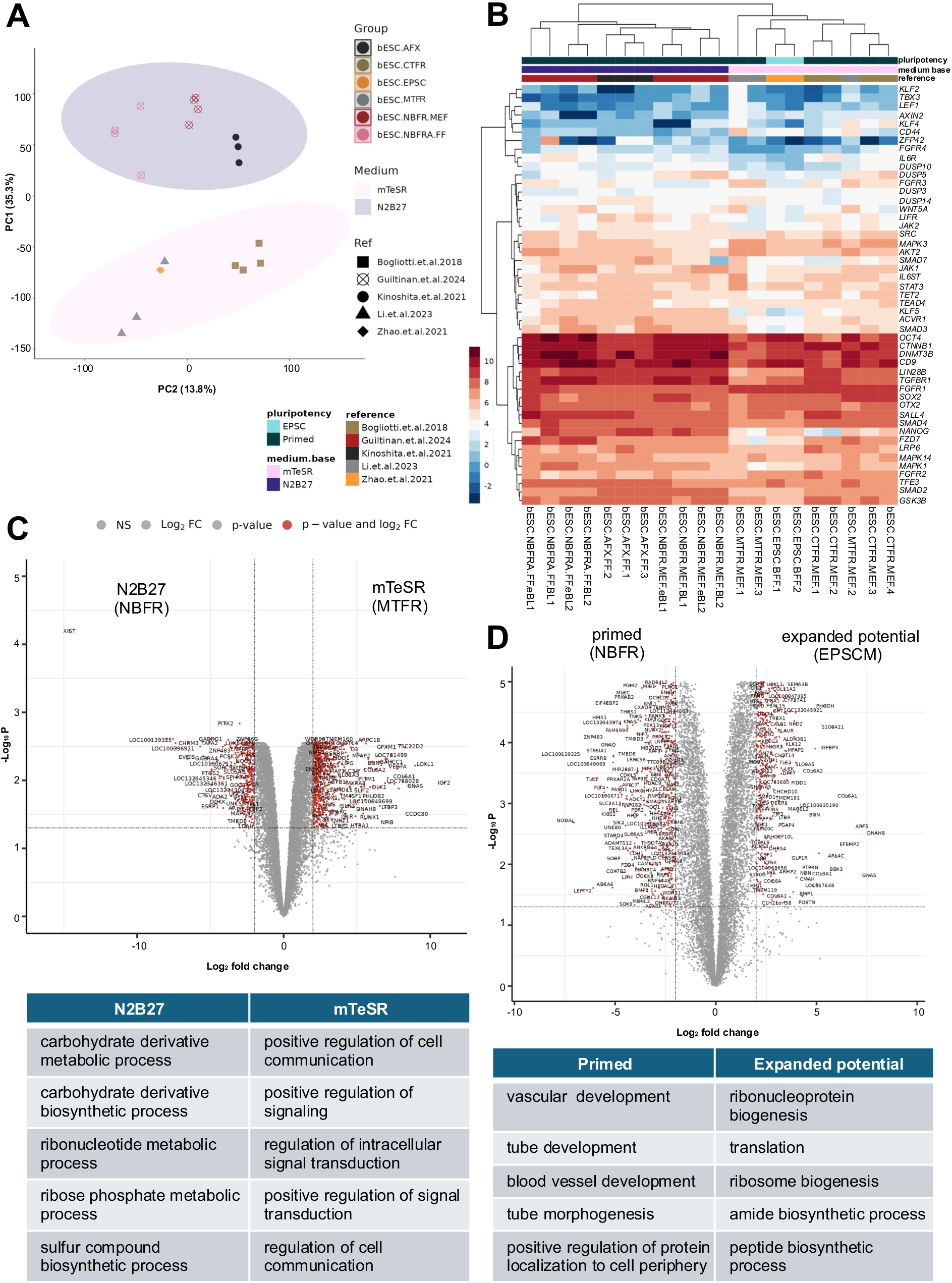
Transcriptomic comparison of bovine embryonic stem cell lines from recent reports. **(A)** Principal component analysis from RNA-seq analysis of bovine ESC lines from this study integrated with publicly-available datasets. **(B)** Heatmap of pluripotency-associated genes among RNA-seq datasets. **(C)** Volcano plot showing differential gene expression based on base medium (mTeSR for MTFR, N2B27 for NBFR) with the same signaling molecules added. Red points indicate differentially expressed genes with log_2_ fold change ≥ 2. Table shows top 5 upregulated biological processes per group based on gene ontology analysis. **(D)** Volcano plot showing differential gene expression between bovine ESCs maintained in media for expanded potential (EPSCM) and primed pluripotency (NBFR). Red points indicate differentially expressed genes with log_2_FC ≥ 2. Table shows top 5 upregulated biological processes per group based on gene ontology analysis.

**Table 1.** Summary of key features of bovine embryonic stem cell lines used for integrative RNA-seq analysis. Green indicates activator, red indicates inhibitor. Abbreviations: BSA = bovine serum albumin, BFF = bovine fetal fibroblast, MEF = mouse embryonic fibroblast, VTN-N = vitronectin, NS = not specified, SE = single-end reads, PE = paired-end reads.

| Source of dataset | Zhao et al., 2021 | Bogliotti et al., 2018 | Li et al., 2023 | Guiltinan et al., 2024 (this report) |  | Kinoshita et al., 2021 |
| --- | --- | --- | --- | --- | --- | --- |
| GEO project ID (PRJNA#) | 532647 | 432598 | 1041850 | *To be updated |  | 723284 |
| Medium-cell name | EPSCM-EPSC | CTFR-ESC | MTFR-ESC | NBFR-ESC |  | AFX-EDSC |
| Pluripotent state | Expanded potential | Primed | Primed | Primed |  | Primed |
| Embryo age (days) | 5-6 | 7-8 | 7-8 | 6-7 |  | 7-9 |
| Base medium | mTeSR | Custom TeSR | mTeSR-E6 | N2B27 |  | N2B27 |
| FGF | N/A | 20 ng/mL FGF2 | 20 ng/mL FGF2 | 20 ng/mL FGF2 |  | 12.5 ng/mL FGF2 |
| WNT | 1 μM CHIR99021, 5 μM IWR-1 /XAV939 | 2.5 μM IWR-1 | 2.5 μM IWR-1 | 2.5 μM IWR-1 | | 2 μM XAV939 |
| TGFβ | 20 ng/mL ACTA | N/A | N/A | N/A | 20 ng/mL ACTA | 20 ng/mL ACTA |
| Other supplements | 0.3 μM WH-4-023, 50 μg/mL Vitamin C, 10 ng/mL LIF | 1% BSA | 1% BSA | 1% BSA | | N/A |
| Matrix | BFF feeders | MEF feeders | MEF feeders | MEF feeders | VTN-N (feeder-free) | Laminin + Fibronectin (feeder-free) |
| Oxygen tension | Normoxia | Normoxia | Normoxia | Normoxia |  | Hypoxia |
| Passage # | NS | 10 | NS | 9 | 23-32 | 14-17 |
| # of lines sequenced | 2 | 4 | 4 | 4 × 2 matrices | | 3 |
| Sequencing platform | Illumina HiSeq 2000 PE | Illumina HiSeq 2500 SE | Illumina NovaSeq 6000 PE | Illumina NovaSeq S4 PE | Illumina HiSeq 4000 PE | Illumina NovaSeq 6000 PE |

With the exception of one report of expanded potential bEPSCs, all other embryo-derived PSCs from cattle have been shown to be in a primed state of pluripotency. With this in mind, we next sought to compare our cell lines, which we determined to be primed based on prior analyses, to bEPSCs. We found that our cells in NBFR had 622 upregulated genes and 500 downregulated genes compared to those in EPSCM (Fig. 4D). Gene ontology analysis revealed that NBFR-bESCs had enriched markers related to advanced development (e.g., vascular, tube, and blood vessel development, tube morphogenesis). On the other hand, bEPSCs in EPSCM had enhanced markers associated with protein biosynthesis and translation (Fig. 4D). These findings further support that NBFR-bESCs are in a primed state of pluripotency in which they are bracing for differentiation into other cell lineages.

## DISCUSSION

Findings from the present study indicate that bovine ESCs can be derived from early and full, non-expanded blastocyst-stage embryos at a similar efficiency (33.3% in this report). Moreover, cell lines have similar phenotypic characteristics and transcriptomic profiles when maintained in the same conditions. It is worth noting that other studies have indicated a significantly higher derivation efficiency for day 8-9 (73.3 and 75%, respectively), as opposed to day 7 (12.5%) bovine embryos (Kinoshita et al., 2021b). One contributing factor to this may be that blastocyst expansion and hatching aid in the process of ICM attachment and outgrowth formation, which is critical for ESC establishment. In our experiment, any embryos that did not attach on their own within one day were manually attached to the feeder cell layer, which likely increased the derivation efficiency for the earlier embryo stages, which were less inclined to attach despite having the ZP enzymatically removed prior to seeding. Moreover, the aforementioned report established lines in feeder-free conditions, whereas this experiment derived lines on MEF feeders, which may have led to less differential advantage for further developed embryos.

The medium used for establishment and maintenance of bESC lines in this study was based on a report using an N2B27 base with supplementation of FGF2, ACTA, and a canonical WNT inhibitor (Soto et al., 2021). While in vitro culture of embryos is typically limited to around 7 days, recent studies have demonstrated the extended culture of bovine and ovine embryos beyond the blastocyst stage with the addition of N2B27 supplements (Ramos-Ibeas et al., 2020; Ramos-Ibeas et al., 2022). Potentially, embryos from both blastocyst stages in this experiment had the opportunity to further develop before their attachment and outgrowth. Moreover, if there were initially differences in epiblast characteristics due to the blastocyst stage, it is possible that all resulting cell lines progressed to a similar point of development by being maintained in primed medium, which stabilizes pluripotent cells resembling the peri-gastrulation epiblast. While this study did not assess the derivation of bESCs from early blastocysts in alternative conditions, future investigations using less advanced embryo stages for derivation of bESCs in media that support naïve and formative pluripotency could be warranted, given that PSCs have yet to be stabilized in these conditions from full-expanded bovine blastocysts.

Interestingly, we generated a high proportion (85%) of female bESC lines in this study. Previous reports have suggested that male bovine embryos may develop more rapidly than female embryos during the pre-implantation period (Avery et al., 1991; Carvalho et al., 1996). Our experimental design tended toward selecting less developmentally-advanced blastocysts within each timepoint (i.e., those that were still at the early blastocyst stage at day 6 and those that had not yet expanded at day 7). This consideration could possibly be used in future studies where it would be advantageous to enrich for a specific sex during cell line establishment. Further experiments with a greater number of embryos will be necessary to determine whether this trend is repeatable.

Previous studies have demonstrated differential FGF requirements depending on pluripotent state, where primed cells require FGF2 supplementation, naïve cells are destabilized by it, and formative PSCs are stable with no to moderate exogenous FGF2 (Kinoshita et al., 2021a; Smith, 2017; Yu et al., 2021). Notably, the only reported instance of bovine PSC culture without FGF2 added to the medium is bEPSCs (Zhao et al., 2021).

Our experiments demonstrate short-term stability of bESCs at concentrations as low as 5 ng/mL of FGF2, which warrants further study over long-term culture and to explore the effects of FGF2 dose on characteristics of pluripotency. The present analysis also pointed to base medium as a source of variation for global transcriptome. While primed bESCs can be stabilized in both mTeSR and N2B27 bases, bEPSCs could only be established in an mTeSR base (Zhao et al., 2021). This was distinct from porcine and human EDSCs, which could be maintained in an N2B27 base with the same cytokines (Gao et al., 2019). This indicates that alternative pluripotent states from primed pluripotency may rely upon the presence of other undetermined factors that affect cellular status, such as metabolic factors.

Aside from differences in media composition, recent reports have described the derivation of PSCs from bovine embryos with varying environmental factors, including alternative matrices (e.g., feeder vs. feeder-free culture; laminin and fibronectin vs. VTN-N; bovine fetal fibroblasts vs. MEFs), and oxygen tension (e.g., normoxia vs. hypoxia; Kinoshita et al., 2021b; Shirasawa et al., 2024; Zhao et al., 2021). However, the effects of these changes on cell line characteristics have not been directly investigated. In addition to the presence and concentration of specific signaling factors in the medium, assessing the effects of environmental factors individually on bESC pluripotency and self-renewal could shed light on alternative sources of variation in the functional capabilities of ESC lines. Collectively, these advances could be combined to determine the optimal conditions for bESC culture for a given application and uncover the determinants of the pluripotency status of cells in vitro.

In all, these findings broaden the potential source of bESC lines to embryos as early as the early blastocyst stage. This could be useful in cases where access to embryos is limited – such as for high genetic merit individuals – or to enable synchronized derivation of ESC lines from a range of blastocyst stages on the same day. Moreover, we have found that the embryo stage from which ESC lines are derived does not have long-term consequences on developmental potential or pluripotent state when cell lines are maintained in the same conditions. Comparative analysis of bovine PSC lines from recent reports shed light on factors that affect the global transcriptome of cell lines, including the base medium and cytokine combinations. Future experiments could address whether earlier staged embryos may be permissive to ESC line establishment in culture conditions that would stabilize a less developmentally-advanced phenotype as compared to later embryonic stages. In addition, experiments extrapolating the effect of specific chemical factors of the medium, as well as environmental factors, will be crucial to dissect how the pluripotent state of ESC lines is determined and specifically which factors can support the establishment and maintenance of bESCs in naïve or formative pluripotency.

## MATERIALS AND METHODS

All reagents and consumables were purchased from Thermo Fisher Scientific unless otherwise specified.

### Derivation and culture of bovine embryonic stem cells

Bovine (*Bos taurus*) embryos were produced in vitro using abattoir-derived ovaries following manufacturer protocols (IVF Biosciences, Falmouth, UK). Quality 1 embryos were staged as early (eBL; stage 5) or full blastocysts (BL; stage 6) according to guidelines from the International Embryo Technology Society Manual (Robertson et al., 1998). Derivation of ESC lines from twelve embryos per blastocyst stage was performed as previously described (Soto et al., 2021), with slight modifications. Upon harvest, embryos were treated with 2 mg/mL Pronase (10165921001, Sigma) for 2-3 min to remove the zona pellucida (ZP), washed several times in SOF-HEPES medium, and plated individually in NBFR medium (N2B27 medium [1:1 DMEM/F12 (11320-033, Gibco) and Neurobasal (21103-049, Gibco) media, 0.5% v/v N-2 supplement (17502-048, Gibco), 1% v/v B-27 supplement (17504-044, Gibco), 2 mM MEM Non-Essential Amino Acid solution (M7145, Sigma), 1% v/v GlutaMAX supplement (35050-061, Gibco), 0.1 mM 2-mercaptoethanol (M6250, Sigma), 100 U/mL Penicillin, and 100 μg/mL Streptomycin (15140-122, Gibco)] supplemented with 1% bovine serum albumin (0219989950, MP Biomedicals), 20 ng/mL fibroblast growth factor 2 (FGF2; 100-18B, PeproTech), and 2.5 μM of the tankyrase inhibitor IWR-1 (I0161, Sigma)) in wells of a 48-well plate that were pre-seeded with mouse embryonic fibroblast (MEF) feeders (A34180, Gibco) on 0.1% gelatin. During the first week of derivation, media was also supplemented with 10 µM Rho Kinase inhibitor Y27632 (ROCKi; ALX-270-333, Enzo) and antibiotic/antimycotic solution (100 U/mL Penicillin, 100 μg/mL Streptomycin, 0.25 μg/mL Amphotericin B; 20004, JR Scientific) rather than antibiotic alone. Any embryos that had not attached after 24 h of culture were manually attached using a 23G needle. Outgrowths were cultured at 37 °C, 5% CO_2_ and normoxia with daily media changes and passaged following single-cell dissociation using TrypLE Express onto fresh MEFs at a 1:1-2 split ratio with 10 μM ROCKi every 7 days.

When ESC derivation was successful, colonies were observed by passage 4. Once established, bESCs were passaged at a 1:5-10 split ratio every 3-4 days and maintained on MEFs or adapted to feeder-free culture on recombinant, truncated vitronectin (VTN-N; A14700, Gibco) in 4-or 6-well plates. For feeder-free and advanced-passage culture, NBFR medium was additionally supplemented with 20 ng/mL activin A (ACTA). Established bESCs were cultured for at least 30 passages with routine testing to confirm that cultures were free of mycoplasma (MP0035, Sigma). Cell morphology was checked daily using a digital inverted microscope (EVOS M5000, Thermo Fisher). Cell counting and viability assessment were performed using an automated cell counter (Countess 3, Invitrogen) and 0.4% Trypan blue cell dye.

### Immunolocalization analysis

Adherent bESC cultures that had been grown to 70-80% confluency were fixed in 4% paraformaldehyde for 10 min at room temperature (RT), and then rinsed and stored in phosphate-buffered saline (PBS) at 4 °C until staining. Cells were permeabilized in 1% Triton X-100 in PBS for 10 min, blocked with 0.3% Triton X-100 and 3% normal donkey serum in PBS for 1 h at RT. Washes were done with 0.3% Triton-X-100 in PBS (PBST), and antibody mixes were prepared in blocking buffer. Cells were incubated with primary antibody mixes at RT for 1 h or at 4 °C overnight. Secondary antibody incubation was done for 1 h at RT and protected from light. Hoechst 33342 (561908, BD Biosciences; 1:1000) in PBST was used for nuclear counterstaining, by incubating for 15 min at RT in the dark. Pluripotency of ESC lines was confirmed by homogenous co-expression of the core pluripotency markers OCT4, SOX2, and NANOG. Epigenetic status was assessed by expression of H3K27me3 and H3K9me2. Antibody dilutions and catalog numbers are listed in Table S1. Cells were imaged using a fluorescent microscope equipped with a high content imaging system (ImageXpressMicro, Molecular Devices). Image processing was performed with the Meta ImageXpress software (v6.7.2.290, Molecular Devices) and ImageJ2 software (v2.9.0/1.53t).

### Sex determination

Sex of bESC lines was determined by polymerase chain reaction (PCR) of the DEAD box helicase 3 gene (*DDX3*), as previously described (Soto et al., 2021). Extraction of DNA was performed using the DNeasy Blood and Tissue Kit (69504, Qiagen) following the manufacturer’s protocol, and quantified using a NanoDrop 2000C Spectrophotometer. Amplification of DNA was performed using GoTaq HotStart Polymerase (M5001, Promega) according to manufacturer’s instructions. Amplicons were visualized by agarose gel electrophoresis with SYBR Safe DNA stain (S33102, Invitrogen) and discrimination between X and Y chromosomes was determined based on sex-specific differences in amplicon size (Gokulakrishnan et al., 2012). Genomic DNA from adult bovine testes and ovaries were used as controls. Primer sequences for *DDX3* amplification and product sizes are shown in Table S2.

### Karyotype analysis

Cell cultures at 70-80% confluency were incubated for 1 h at 37 °C in 10 μg/mL Karyomax Colcemid solution in HBSS at a working concentration of 0.1 μg/mL. After this period, media was discarded, wells were rinsed twice with PBS, and cells were detached from the plate using TrypLE Express reagent. Cells were collected into a 15 mL conical tube and centrifuged at 300 x g for 2 min. Cell pellets were slowly and gently resuspended into a single-cell suspension in pre-warmed (37 °C) hypotonic solution (0.56% KCl in deionized H2O) using Pasteur pipettes with bulbs. The single-cell KCl solution was incubated at 37 °C for 25 min then spun down at 300 x g for 5 min, for a total time of 30 min in KCl solution. The hypotonic solution was drawn off and the cells were fixed using 3:1 Methanol to Glacial Acetic Acid at RT for 15 min. The fixative was then drawn off, new 3:1 fixative was added, and cells were gently re-suspended until uniformly mixed without any clumps. Fixed cells were stored at 4 °C overnight. The following day, the cell suspension was pelleted at 300 x g for 5 min and resuspended in freshly-made 3:1 fixative. Using a glass Pasteur pipet, the cell suspension was drawn up and dropped onto a microscope slide from 6-8 inches above and then allowed to air dry. Once dry, the slides were stained using 3% Giemsa/PBS stain solution (LC148407, LabChem) for 3 min, and then rinsed two times in DI water. After letting air dry for 5 min, metaphase spreads were imaged at 45x on a phase microscope (AO One-Fifty, American Optical). Four cell lines (n = 2/blastocyst stage) were analyzed through 30 passages.

### Teratoma formation

Bovine ESCs from each blastocyst stage (n = 1/line) at passage 12-14 were grown on MEFs to 70% confluency and collected as they would be for passaging. Cell pellets were resuspended in PBS, counted, and the appropriate volume was resuspended in PBS and cold 60% growth-factor reduced Matrigel (CLS356230, Corning) at a 1:1 ratio, such that the final solution was 30% Matrigel with 10,000 cells/μL. Cells suspensions were transported on ice to the UC Davis Stem Cell Core Facility. One million cells (100 μL solution) per cell line were injected into the upper hind leg of an immunodeficient mouse. After twelve weeks, mice were sacrificed and the leg that had been injected was fixed in formalin. Samples were transferred into 70% ethanol after 48 h. Sectioning and hematoxylin and eosin (H&E) staining were performed by the Comparative Pathology Laboratory at UC Davis. Assessment for presence of the three embryonic germ layers was assessed by a board-certified veterinary pathologist. All animal work was performed under the approval of the Institutional Animal Care and Use Committee at UC Davis (#23335).

### RNA isolation

Isolation of RNA was performed using the Purelink RNA Mini Kit (12183018A, Invitrogen) or RNeasy Plus Mini Kit (74143, Qiagen) following manufacturer protocols with DNase treatment (12185010, Invitrogen). Samples that were snap frozen and stored at –80 °C were processed using the kit-provided lysis buffers, whereas those stored in TRIzol reagent underwent phase separation in chloroform following manufacturer instructions before proceeding with the kit. Quantity and quality of RNA was assessed using a Nanodrop 2000C Spectrophotometer. Within a given experiment, RNA concentration was normalized by diluting samples in nuclease-free water.

### Reverse transcription-quantitative polymerase chain reaction (RT-qPCR)

Reverse transcription of RNA products into cDNA was done using the RevertAid First Strand cDNA Synthesis Kit (K1622, Thermo Scientific). Then, qPCR of cDNA products was performed using SsoAdvance Universal SYBR Green Supermix (1725271, BioRad) reagent for 40 cycles on the CFX96 Touch Deep Well Real-Time PCR Detection System (BioRad) following manufacturer instructions. The gene H2A.2 was used as a reference marker for delta-cycle threshold (ΔCT) calculation. The ΔΔCT value was calculated by subtraction the ΔCT of a given treatment from the control condition (i.e., 20 ng/mL FGF2 for this experiment). Fold change was calculated as 2^-ΔΔCT^. Primer sequences used for RT-qPCR analysis are listed in Table S2.

### RNA-sequencing analysis

Sequencing of RNA from NBFR-bESC lines cultured on MEFs (n = 2 lines/embryo stage) at passage 9 was performed at the UC Davis DNA Technology Core Facility using PolyA enrichment (NovaSeq S4 PE150) with 25 million strand-specific paired reads. Sequenced reads were aligned to the bovine genome (NCBI ARS-UCD 2.0) using STAR and raw counts matrices were prepared using FeatureCounts. Preliminary quality control analysis was performed using FastQC and sequencing adaptors were trimmed from the raw data using Trimmomatic. The read numbers mapping to each gene were calculated using the Feature Counts. Data analysis was performed using R software (v4.2). One sample of pure MEFs was sequenced and any reads with alignment to the bovine genome were removed from sample count matrices before analysis to account for presence of MEFs in bESC samples. Raw counts were converted into counts per million using EdgeR, then genes with low counts were removed and the TMM (i.e., Trimmed Mean of M-values) function was applied to standardize the counting data of the reads. EdgeR was used to analyze the differentially expressed genes (DEGs). P-values were adjusted for the false discovery rate. Genes considered to be DEGs had threshold values of adjusted p-value < 0.05 and fold change ≥ 2. We also integrated RNA-seq data that we generated from the same bESC lines following feeder-free transition onto VTN-N and advanced culture to passage 23-32 (HiSeq 4000 PE150), as well as published data of epiblast samples from bovine embryos at day 7, 14, and 17 of embryonic development (Bernardo et al., 2018). Moreover, we performed a comparative analysis of our NBFR-ESC lines with publicly-available RNA-seq datasets including primed CTFR-bESCs (Bogliotti et al., 2018; Li et al., 2023) and AFX-bEDSCs (Kinoshita et al., 2021b), as well as expanded potential EPSCM-bEPSCs (Zhao et al., 2021). Raw count files were downloaded and analyzed following the same pipeline described previously. Gene ontology analysis of enriched biological processes based on differentially expressed genes was performed using the GAGE package in R.

**Figure S1.**
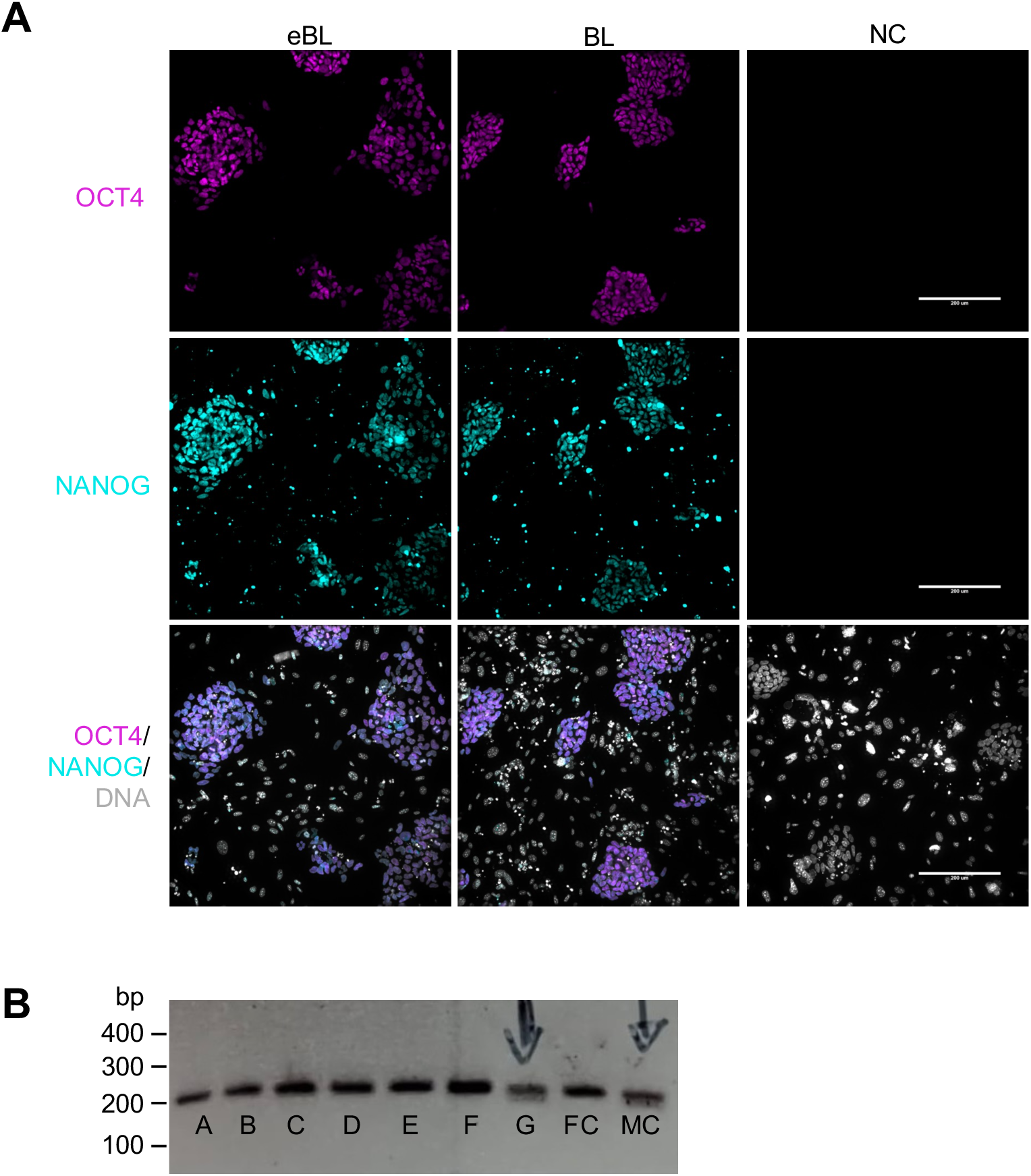
Additional characterization of bovine ESC lines from this study. **(A)** Representative immunofluorescence images of bESCs derived from early (eBL) and full (BL) blastocysts on MEF feeders at passage 16. Magenta indicates OCT4; cyan indicates NANOG; gray indicates DNA. NC is negative control, with no primary antibody added. Scale bar, 200 μm. **(B)** Sexing results of bESC lines visualized by agarose gel electrophoresis. A-G represent tested cell lines from this experiment. MC is male control, FC is female control. Arrows indicate samples determined as male, based on two bands.

**Table S1.** Antibodies used for immunolocalization analysis. Primary antibodies were reconstituted to the concentration specified by manufacturer instructions.

| Primary antibodies | Manufacturer | Catalog number | Dilution |
| --- | --- | --- | --- |
| Goat anti-OCT3/4 | R&D Systems | AF1759 | 1:100 |
| Rabbit anti-SOX2 | BioGenex | NU833-5UC | 1:300 |
| Rabbit anti-NANOG | Invitrogen | 500-P236 | 1:200 |
| Rabbit anti-H3K27me3 | Millipore | ABE44 | 1:500 |
| Mouse anti-H3K9me2 | Abcam | AB1220 | 1:500 |
| Secondary antibodies | Manufacturer | Catalog number | Dilution |
| Donkey anti-goat AlexaFluor 568 | Invitrogen | A11057 | 1:500 |
| Donkey anti-rabbit AlexaFluor 488 | Invitrogen | A21206 | 1:500 |
| Donkey anti-mouse AlexaFluor 546 | Invitrogen | A10036 | 1:500 |
| Nuclear dye | Manufacturer | Catalog number | Dilution |
| Hoechst 33342 | BD Biosciences | BDB561908 | 1:1000 |

**Table S2.**
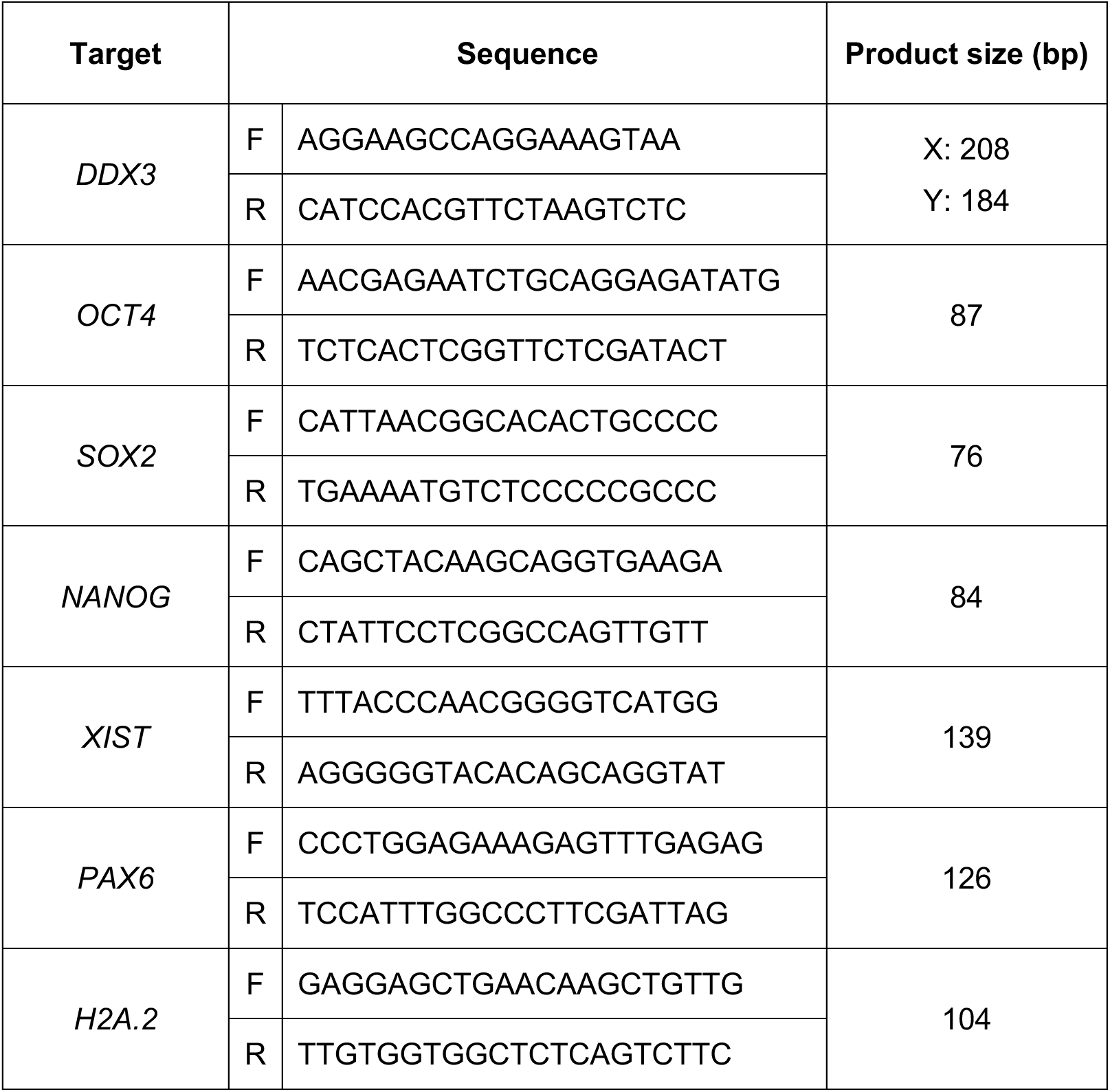
Primer sequences for genes analyzed by polymerase chain reaction. Tm = melt temperature, F = forward and R = reverse primer sequences. X and Y indicate female and male sex chromosomes, respectively.

**Table S3.** Gene ontology analysis of enriched biological processes of bovine ESCs in various conditions. Enriched pathways based on differential gene expression of bovine ESCs cultured in N2B27 base (NBFR) vs. mTeSR-E6 base (MTFR) on top and bovine ESCs cultured in primed medium (NBFR) vs. expanded potential medium (EPSCM) on bottom.

| **MTFR vs. NBFR** | p.geomean | stat.mean | p.val | q.val | set.size | exp1 |
| --- | --- | --- | --- | --- | --- | --- |
| GO:1902531 regulation of intracellular signal transduction | 0.0018 | 2.9294 | 0.0018 | 0.6083 | 207 | 0.0018 |
| GO:0010647 positive regulation of cell communication | 0.0019 | 2.9158 | 0.0019 | 0.6083 | 187 | 0.0019 |
| GO:0023056 positive regulation of signaling | 0.0020 | 2.8908 | 0.0020 | 0.6083 | 188 | 0.0020 |
| GO:0009967 positive regulation of signal transduction | 0.0026 | 2.8179 | 0.0026 | 0.6083 | 171 | 0.0026 |
| GO:0048583 regulation of response to stimulus | 0.0029 | 2.7647 | 0.0029 | 0.6083 | 448 | 0.0029 |
| GO:0010646 regulation of cell communication | 0.0032 | 2.7363 | 0.0032 | 0.6083 | 372 | 0.0032 |
| GO:0009966 regulation of signal transduction | 0.0033 | 2.7207 | 0.0033 | 0.6083 | 333 | 0.0033 |
| GO:0023051 regulation of signaling | 0.0034 | 2.7098 | 0.0034 | 0.6083 | 373 | 0.0034 |
| GO:1902533 positive regulation of intracellular signal transduction | 0.0041 | 2.6694 | 0.0041 | 0.6083 | 116 | 0.0041 |
| GO:0048513 animal organ development | 0.0045 | 2.6229 | 0.0045 | 0.6083 | 271 | 0.0045 |
| GO:0035556 intracellular signal transduction | 0.0049 | 2.5903 | 0.0049 | 0.6083 | 299 | 0.0049 |
| GO:0008285 negative regulation of cell population proliferation | 0.0056 | 2.5772 | 0.0056 | 0.6083 | 74 | 0.0056 |
| GO:0061061 muscle structure development | 0.0066 | 2.5198 | 0.0066 | 0.6083 | 65 | 0.0066 |
| GO:0048584 positive regulation of response to stimulus | 0.0071 | 2.4611 | 0.0071 | 0.6083 | 252 | 0.0071 |
| GO:0008283 cell population proliferation | 0.0101 | 2.3283 | 0.0101 | 0.6083 | 315 | 0.0101 |
| **NBFR vs. MTFR** | p.geomean | stat.mean | p.val | q.val | set.size | exp1 |
| GO:0044272 sulfur compound biosynthetic process | 0.0117 | -2.3790 | 0.0117 | 0.9968 | 19 | 0.0117 |
| GO:1901135 carbohydrate derivative metabolic process | 0.0135 | -2.2234 | 0.0135 | 0.9968 | 127 | 0.0135 |
| GO:1901137 carbohydrate derivative biosynthetic process | 0.0185 | -2.1066 | 0.0185 | 0.9968 | 72 | 0.0185 |
| GO:0019693 ribose phosphate metabolic process | 0.0313 | -1.8827 | 0.0313 | 0.9968 | 53 | 0.0313 |
| GO:0009101 glycoprotein biosynthetic process | 0.0330 | -1.8958 | 0.0330 | 0.9968 | 21 | 0.0330 |
| GO:0009259 ribonucleotide metabolic process | 0.0351 | -1.8297 | 0.0351 | 0.9968 | 52 | 0.0351 |
| GO:0033865 nucleoside bisphosphate metabolic process | 0.0407 | -1.8210 | 0.0407 | 0.9968 | 13 | 0.0407 |
| GO:0033875 ribonucleoside bisphosphate metabolic process | 0.0407 | -1.8210 | 0.0407 | 0.9968 | 13 | 0.0407 |
| GO:0034032 purine nucleoside bisphosphate metabolic process | 0.0407 | -1.8210 | 0.0407 | 0.9968 | 13 | 0.0407 |
| GO:0032392 DNA geometric change | 0.0422 | -1.7763 | 0.0422 | 0.9968 | 21 | 0.0422 |
| GO:0071103 DNA conformation change | 0.0422 | -1.7763 | 0.0422 | 0.9968 | 21 | 0.0422 |
| GO:0009156 ribonucleoside monophosphate biosynthetic process | 0.0456 | -1.8318 | 0.0456 | 0.9968 | 10 | 0.0456 |
| GO:0070085 glycosylation | 0.0467 | -1.7323 | 0.0467 | 0.9968 | 19 | 0.0467 |
| GO:0006261 DNA-templated DNA replication | 0.0468 | -1.7173 | 0.0468 | 0.9968 | 26 | 0.0468 |
| **EPSCM vs. NBFR** | p.geomean | stat.mean | p.val | q.val | set.size | exp1 |
| GO:0022613 ribonucleoprotein complex biogenesis | 0.0001 | 3.7140 | 0.0001 | 0.1959 | 85 | 0.0001 |
| GO:0006412 translation | 0.0006 | 3.2856 | 0.0006 | 0.2907 | 122 | 0.0006 |
| GO:0042254 ribosome biogenesis | 0.0006 | 3.3161 | 0.0006 | 0.2907 | 55 | 0.0006 |
| GO:0043604 amide biosynthetic process | 0.0009 | 3.1403 | 0.0009 | 0.3178 | 138 | 0.0009 |
| GO:0043043 peptide biosynthetic process | 0.0012 | 3.0584 | 0.0012 | 0.3204 | 127 | 0.0012 |
| GO:0044085 cellular component biogenesis | 0.0014 | 2.9958 | 0.0014 | 0.3204 | 354 | 0.0014 |
| GO:0043933 protein-containing complex organization | 0.0025 | 2.8144 | 0.0025 | 0.4401 | 261 | 0.0025 |
| GO:0006518 peptide metabolic process | 0.0027 | 2.8041 | 0.0027 | 0.4401 | 144 | 0.0027 |
| GO:0065003 protein-containing complex assembly | 0.0029 | 2.7730 | 0.0029 | 0.4401 | 191 | 0.0029 |
| GO:0045861 negative regulation of proteolysis | 0.0042 | 2.8298 | 0.0042 | 0.5643 | 18 | 0.0042 |
| GO:0044271 cellular nitrogen compound biosynthetic process | 0.0051 | 2.5754 | 0.0051 | 0.6278 | 420 | 0.0051 |
| GO:0007005 mitochondrion organization | 0.0062 | 2.5244 | 0.0062 | 0.6451 | 92 | 0.0062 |
| GO:0008202 steroid metabolic process | 0.0065 | 2.5875 | 0.0065 | 0.6451 | 24 | 0.0065 |
| GO:0022900 electron transport chain | 0.0067 | 2.5923 | 0.0067 | 0.6451 | 22 | 0.0067 |
| GO:0032543 mitochondrial translation | 0.0086 | 2.4859 | 0.0086 | 0.7169 | 25 | 0.0086 |
| **NBFR vs. EPSCM** | p.geomean | stat.mean | p.val | q.val | set.size | exp1 |
| GO:0001944 vasculature development | 0.0237 | -2.0101 | 0.0237 | 0.9843 | 47 | 0.0237 |
| GO:0035295 tube development | 0.0251 | -1.9749 | 0.0251 | 0.9843 | 70 | 0.0251 |
| GO:0001568 blood vessel development | 0.0296 | -1.9111 | 0.0296 | 0.9843 | 46 | 0.0296 |
| GO:0035239 tube morphogenesis | 0.0316 | -1.8778 | 0.0316 | 0.9843 | 54 | 0.0316 |
| GO:1904377 positive regulation of protein localization to cell periphery | 0.0330 | -1.9786 | 0.0330 | 0.9843 | 10 | 0.0330 |
| GO:0072359 circulatory system development | 0.0358 | -1.8158 | 0.0358 | 0.9843 | 72 | 0.0358 |
| GO:0007041 lysosomal transport | 0.0380 | -1.9357 | 0.0380 | 0.9843 | 10 | 0.0380 |
| GO:0007254 JNK cascade | 0.0420 | -1.8096 | 0.0420 | 0.9843 | 12 | 0.0420 |

## Acknowledgements

The authors would like to thank Dr. M. Belen Rabaglino for initial transcriptomic analysis of bESC lines. We also extend our gratitude to Alexys Dailly for assistance with cell culture and experiments.

## Author contributions

Conceptualization: C.G., J.I.C., A.C.D.; Methodology: C.G., R.C.B., J.I.C., J.M.S., A.C.D.; Validation: C.G., R.C.B., R.B.A.; Formal analysis: C.G., R.C.B., A.C.D.; Investigation: C.G., R.C.B., J.I.C., J.M.S.; Data curation: C.G., R.C.B., A.C.D.; Writing – original draft: C.G.; Writing – review & editing: C.G., R.C.B., J.I.C, J.M.S., R.B.A, A.C.D.; Supervision: A.C.D.; Project administration: C.G., A.C.D.; Funding acquisition: C.G., A.C.D.

## Data and resource availability

RNA-sequencing datasets will be publicly accessible via GEO (accession codes will be updated when available).

